# Disruption of vacuolin microdomains in the host *Dictyostelium discoideum* increases resistance to *Mycobacterium marinum*-induced membrane damage and infection

**DOI:** 10.1101/2021.11.16.468763

**Authors:** Cristina Bosmani, Angélique Perret, Florence Leuba, Aurélie Guého, Nabil Hanna, Thierry Soldati

## Abstract

*Mycobacterium tuberculosis* (Mtb), the causative agent of tuberculosis, manipulates the host phagosome maturation pathway to replicate intracellularly. *Mycobacterium marinum*, a closely-related species, and *Dictyostelium discoideum*, a social amoeba and alternative phagocytic host, have been used as models to study host-pathogen interactions occurring during mycobacterial infections. Vacuolins, functional homologues of the mammalian flotillins, organize membrane microdomains and play a role in vesicular trafficking. Various pathogens have been reported to manipulate their membrane association and function. During infection of *D*. *discoideum* with *M*. *marinum*, Vacuolin C was specifically and highly induced and all three vacuolin isoforms were enriched at the mycobacteria-containing-vacuole (MCV). In addition, absence of vacuolins reduced escape from the MCV and conferred resistance to *M*. *marinum* infection. Moreover, ESAT-6, the membrane-disrupting virulence factor of *M*. *marinum*, was less associated with membranes when vacuolins were absent. Together, these results suggest that vacuolins are important host factors that are manipulated by mycobacteria to inflict membrane damage and escape from their compartment.

## INTRODUCTION

Tuberculosis, an infectious disease caused by the pathogen *Mycobacterium tuberculosis* (Mtb), is the 1^st^ cause of death by an infectious disease worldwide and killed 1.4 million people in 2019 (World Health Organization, 2021). Alveolar macrophages are the first line of defense against Mtb, as they take them up by phagocytosis in the lungs to restrict infection. However, Mtb is able to manipulate the phagosomal maturation pathway to eventually establish a replicative niche inside a modified phagosome (Russell, 2001, 2007). *Mycobacterium marinum*, closely related to Mtb, causes a disease similar to tuberculosis in marine and freshwater vertebrates. *M marinum* has been widely used as an experimentally versatile model for Mtb, owing to its faster replication time, easier laboratory manipulation and conserved virulence factors (Stinear et al., 2008; Tobin and Ramakrishnan, 2008).

*Dictyostelium discoideum*, a social amoeba, uses phagocytosis to feed on soil bacteria. The phagosomal maturation pathway is extremely well conserved between *D*. *discoideum* and animal cells (Boulais et al., 2010). After uptake, bacteria are enclosed in a compartment called a phagosome, which matures through a series of fusion and fission events. In the phagosome, bacteria are exposed to acidic pH, proteolytic enzymes, reactive oxygen species (ROS) and toxic levels of metals (Cosson and Lima, 2014; Dunn et al., 2018; Barisch et al., 2018; Hanna et al., 2021). These factors, also conserved between *D*. *discoideum* and mammals, are necessary to achieve killing and digestion of the ingested bacterium. Because of its ease-of-use and conserved phagosomal maturation pathway, *D*. *discoideum* has been widely used as a surrogate macrophage to study host interactions with intracellular pathogens (Cosson and Soldati, 2008; Bozzaro and Eichinger, 2011; Dunn et al., 2018).

The *D*. *discoideum* – *M*. *marinum* system has been previously established as a model to study mycobacterial infections (Solomon et al., 2003; Hagedorn and Soldati, 2007). Like Mtb, after uptake, *M*. *marinum* arrests phagosomal maturation to allow its replication, and eventually escapes from the compartment in an ESX-1-dependent manner (Lewis et al., 2003; Gao et al., 2004; Cardenal-Muñoz et al., 2018). Similarly to Mtb, *M*. *marinum* avoids acidification of the compartment by actively inhibiting association and/or inducing recycling of the vacuolar H^+^-ATPase (v-ATPase; (Wong et al., 2011; Kolonko et al., 2014)). This was shown to require the WASH complex-mediated actin polymerization on the MCV at early time points in both *D*. *discoideum* and macrophages (Kolonko et al., 2014). In addition, the MCV was shown to be almost devoid of cathepsin D and to have a low proteolytic activity (Hagedorn and Soldati, 2007; Cardenal-Muñoz et al., 2017). After establishing its niche, *M*. *marinum* proliferates in the MCV, which becomes more spacious and acquires postlysosomal proteins, such as the predicted copper transporter p80 (Kolonko et al., 2014). *M*. *marinum* eventually escapes from the compartment to access its nutrients and egress from the host cell (Hagedorn and Soldati, 2007; Hagedorn et al., 2009). Throughout the infection course and as early as 1.5 hours post infection (hpi), *M*. *marinum* perforates its compartment thanks to virulence factors, namely the ESAT-6 peptide, secreted through its Type VII secretion system (T7SS) ESX-1 (Cardenal-Muñoz et al., 2017; López-Jiménez et al., 2018). *M*. *marinum* lacking the ESX-1 secretion system (ΔRD1 mutant) is unable to arrest phagosomal maturation and to grow efficiently intracellularly (Hagedorn and Soldati, 2007; Cardenal-Muñoz et al., 2017; López-Jiménez et al., 2018). Thus, the membrane damaging ability of *M*. *marinum* is a prerequisite for efficient replication inside its host.

Flotillin-1 and −2 are highly conserved proteins found in lipid rafts at the plasma membrane and phagosomes (Dermine et al., 2001; Morrow and Parton, 2005; Otto and Nichols, 2011). They insert into the cytosolic leaflet of membranes thanks to their PHB domain (prohibitin homology domain) and acylations at the N-terminus. In addition, coiled-coil regions at the C-terminus allow homo- and heterotetramerization and thus formation of specific membrane microdomains (Morrow et al., 2002; Neumann-Giesen et al., 2004; Solis et al., 2007). Flotillins have been proposed to function as signaling platforms and in membrane trafficking of cargoes (Babuke and Tikkanen, 2007; Stuermer, 2011). Flotillin-1 is present on phagosomes and intracellular pathogens have been shown to interfere with its association with their vacuole. For instance, flotillin-1 is present at the *Brucella abortus*-containing compartment as well as the *Chlamydia pneumoniae* inclusion throughout infection (Arellano-Reynoso et al., 2005; Korhonen et al., 2012). In addition, *Anaplasma phagocytophilum* inclusions are also enriched with flotillins, and it was proposed that flotillins participate in trafficking of free cholesterol to allow replication of *A*. *phagocytophilum* (Xiong et al., 2019). Recently, flotillins were found to be associated with the *Pseudomonas aeruginosa* lectin LecA, and to be required for bacteria entry into the host cell, together with saturated long-chain fatty acids and the GPI-anchored protein CD59 (Brandel et al., 2021). On the other hand, the *Leishmania* parasite actively depletes flotillin-1 from its compartment by preventing phagosome-lysosome fusion (Dermine et al., 2001). Together, these results point to an important role of specific flotillin-enriched microdomains in host-pathogen interactions.

We, and others, have previously shown that the three *D*. *discoideum* vacuolins (VacA, B and C) are flotillin homologues (Jenne et al., 1998; Wienke et al., 2006; Bosmani et al., 2020). They share a similar protein structure, are able to oligomerize and behave as integral membrane proteins. All three vacuolins are found associated with membranes of different endocytic compartments, and accumulate at postlysosomes (Bosmani et al., 2020). Previously, VacB was shown to be important for *M*. *marinum* infection, as its absence impaired intracellular growth as well as allowed the v-ATPase to accumulate at the MCV (Hagedorn and Soldati, 2007). We have since then shown that the knock-out (KO) mutant previously used is in fact a multiple vacuolin KO, and have thus generated new vacuolin mutants in the Ax2(Ka) *D*. *discoideum* background (Bosmani et al., 2020). We subsequently showed that absence of vacuolins greatly affects uptake of various types of particles, including *M*. *marinum*, via reduced expression of Myosin VII and impaired recognition and adhesion to particles. In addition, in absence of vacuolins, phagosomes had an earlier reneutralization phase; delivery and/or retrieval of certain lysosomal enzymes was affected, without any impact on bacteria killing (Bosmani et al., 2020).

In light of the roles recently proposed for vacuolins in *D*. *discoideum*, we sought to better dissect their link with the MCV and their role during *M*. *marinum* infection. We show here that VacC is highly induced upon *M*. *marinum* infection and that vacuolins associate with the MCV as early as 1 hpi. Moreover, we confirm that absence of vacuolins confers resistance to infection, and propose that vacuolins are required for *M*. *marinum* to efficiently damage its compartment and escape to the cytosol where it continues to replicate.

## RESULTS

### Vacuolin C is specifically induced upon *M*. *marinum* infection

To test whether mycobacteria manipulate the expression of the three vacuolin isoforms during the course of the infection, the transcriptomic response of wild-type (wt) Ax2(Ka) cells infected with GFP-expressing *M*. *marinum* was analyzed as previously described (Hanna et al., 2019). The *vacC* gene, which is normally poorly expressed in vegetative cells (dictyExpress, (Stajdohar et al., 2017)), was consistently and significantly induced throughout the 48 hours of infection (**Fig. 1A**). Interestingly, *vacC* was highly induced as early as 1 hour post infection (hpi). On the other hand, *vacB* was only highly induced at 1 hpi, while expression of *vacA* was mostly upregulated at later time points (24-48 hpi). We wondered whether induction of *vacC* was specific to infection with mycobacteria, or a general response to bacteria phagocytosis. We analyzed the transcriptomic response of Ax2(Ka) and DH1 wt cells in contact for 4 hours with different Gram-positive and -negative bacteria, as well as different mycobacterial strains (**Fig. 1B**, (Lamrabet et al., 2020)). The *vacC* gene was not significantly expressed when cells were in contact with the Gram-positive or -negative strains tested, but consistently highly induced with mycobacteria (**Fig. 1B**). Moreover, the expression of *vacC* correlated with the pathogenicity of the mycobacterial strain, with the highest induction observed with wt *M*. *marinum*, and the lowest with *M*. *smegmatis*, a non-pathogenic mycobacterium. RNAseq results were overall confirmed by qRT-PCR (**Fig. 1C**). Note that, as *vacC* is a poorly expressed gene in basal conditions, the absolute fold change values reported here are higher than observed by RNAseq, due to the lower sensitivity of qRT-PCR. Contrary to the consistent upregulation of *vacC* with wt *M*. *marinum*, infection with the ΔRD1 mutant and *M*. *smegmatis* induced *vacC* expression only at 1 hpi. At later time points, expression of *vacC* was mostly similar to levels observed in non-infected cells. To test whether the VacC protein was correspondingly upregulated upon *M*. *marinum* infection, the endogenous protein levels of each vacuolin was assessed by western blot using the previously described chromosomal Vac-GFP knock-in strains (Vac-GFP KI, (Bosmani et al., 2020). The VacC protein accumulated from 6 hpi and significantly at 24 hpi, compared to mock-infected cells (NI, **Fig. 1D-E**), whereas no significant upregulation was observed for VacA or VacB. In addition, VacC was only highly induced by infection with wt *M*. *marinum*, but not the ΔRD1 mutant **(Fig. 1D-E**). Because we analyzed the level of expression in a heterogenous pool of cells (i.e. including infected and non-infected cells), the exact fold change of VacC was highly dependent on the percentage of infected cells. To confirm whether only infected cells showed an upregulation of VacC, infected Vac-GFP KI cells were imaged by high content microscopy (**Fig. 1F-G**), and levels of GFP intensity were measured in infected versus non-infected cells present in the same pool. The expression of VacC was consistently higher in cells infected with wt *M*. *marinum*, when compared to their non-infected neighbors. No difference in the VacC-GFP level was observed when cells were infected with the ΔRD1 mutant. These results show that VacC is a specific host reporter for the infection with *M*. *marinum*, and its high expression is possibly triggered by ESX-1 dependent damages.

**Figure 1:**
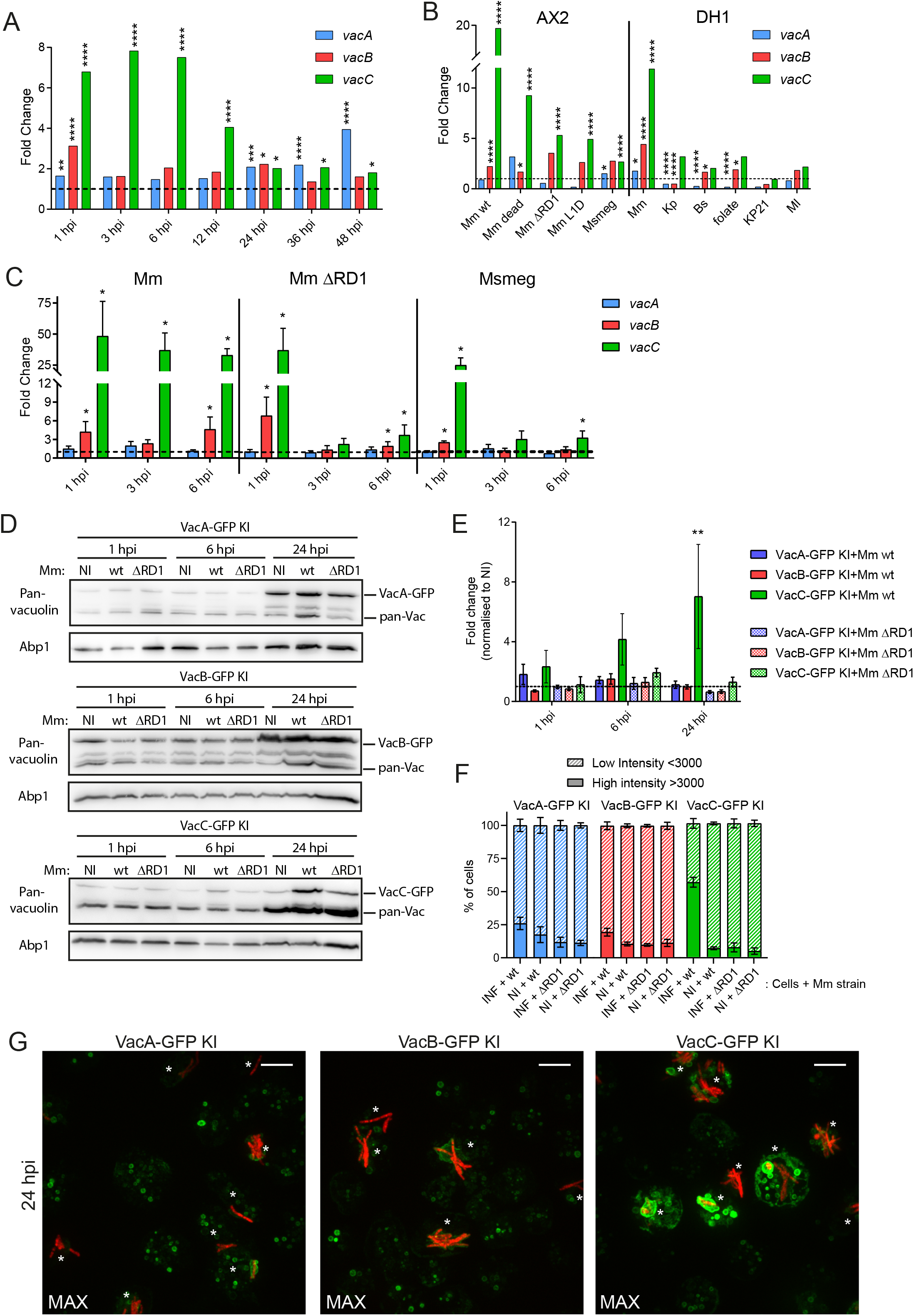
Vacuolin C is specifically induced upon *M*. *marinum* infection. **A.** RNA-sequencing of wt cells infected with GFP-expressing *M*. *marinum* FACSorted at different times post infection (hpi). The population of GFP-positive cells were FACSorted prior to RNA extraction. Fold change RNA levels normalized to mock infected cells and compared to mock-infected cells (dashed line, N=3, *p≤0.05, **p≤0.01, ***p≤0.001, ****p≤0.0001). **B.** RNA-sequencing of AX2 or DH1 *D*. *discoideum* wt cells in contact with the indicated bacterial strains for 4 hours, normalized and compared to mock cells (dashed line). The population of GFP-positive cells were FACSorted prior to RNA extraction (N=3, *p≤0.05, ****p≤0.0001). Mm: *M*. *marinum*, Msmeg: *M*. *smegmatis*, Kp: *Klebiella pneumoniae*, Bs: *Bacillus subtilis*, Ml: *Micrococcus luteus*. **C.** Quantitative RT-PCR of wt cells infected with wt or ΔRD1 *M*. *marinum*, or *M*. *smegmatis*. RNA levels normalized to GAPDH and to mock-infected cells. Cells were not sorted, the samples are heterogenous and contain both infected and non-infected cells (dashed line, mean ± s.e.m., N=4, *p≤0.05, Mann-Whitney test). **D.** Lysate of non-sorted cell lines infected with wt or ΔRD1 *M*. *marinum*, or mock-infected (NI), at different hpi, immunoblotted with the indicated antibodies, representative blot. **E.** Quantification of bands of Vac-GFPs in D., normalized to Abp1 and NI cells (dashed line, N≥3, **p≤0.01, one-way ANOVA). **G.** Representative Max projections of indicated cell lines infected with wt *M*. *marinum* expressing mCherry at 24 hpi, the same settings were used to image all cell lines. *, infected cells; scale bar, 10 μm. **F.** Percentage of infected or non-infected cells of the indicated cell lines with a low or high intensity GFP signal at 24 hpi (mean ± s.e.m., N=2, n≥150 cells).

### Vacuolins are present at the *M*. *marinum*-containing vacuole

It was previously shown that vacuolins decorate the *M*. *marinum* containing vacuole (MCV) throughout the infection course starting as early as 6 hpi (Hagedorn and Soldati, 2007), however, the anti-vacuolin (pan-vacuolin) antibody used recognizes all three isoforms (Bosmani et al., 2020). We therefore wanted to better dissect which vacuolin is mostly associated with the MCV and the dynamics of their recruitment at earlier time-points, given our RNAseq data (Fig.1). Vac-GFP KI cells were infected with mCherry-expressing *M*. *marinum*, and the dynamic of each vacuolin’s recruitment was followed by live microscopy (**Fig. 2A-B**). Each vacuolin was present at the MCV as early as 1 hpi, with an average of 60% of MCVs being decorated with one of the vacuolins the first day of infection (**Fig. 2B**). Interestingly, we observed that the way vacuolins associated with the MCV membrane was different over time. In fact, at early time points, vacuolins exhibited a patchy distribution at the MCV, with only certain regions of the MCV harboring vacuolins (**Fig. 2A and C**). At 24 hpi, on the other hand, the whole MCV membrane was strongly enriched with each vacuolin, with about 75% of vacuolin-positive MCVs having a “solid” vacuolin coat (**Fig. 2C**). These results were confirmed using antibodies against the endogenous VacA and VacB, which were also found to be highly accumulated at the MCV membrane at 24 hpi (**Fig. S1**). Furthermore, at that stage of infection, the damage inflicted by *M*. *marinum* to its MCV became macroscopically obvious (**Fig. 2A**, arrowheads), and only about 30-40% of MCVs had an intact vacuolin-coat (**Fig. 2D**). Overall, these results show that all three vacuolins are present at the MCV, as early as 1 hpi, and gradually accumulate at the MCV membrane until a solid “vacuolin-coat” completely envelops the compartment.

**Figure 2:**
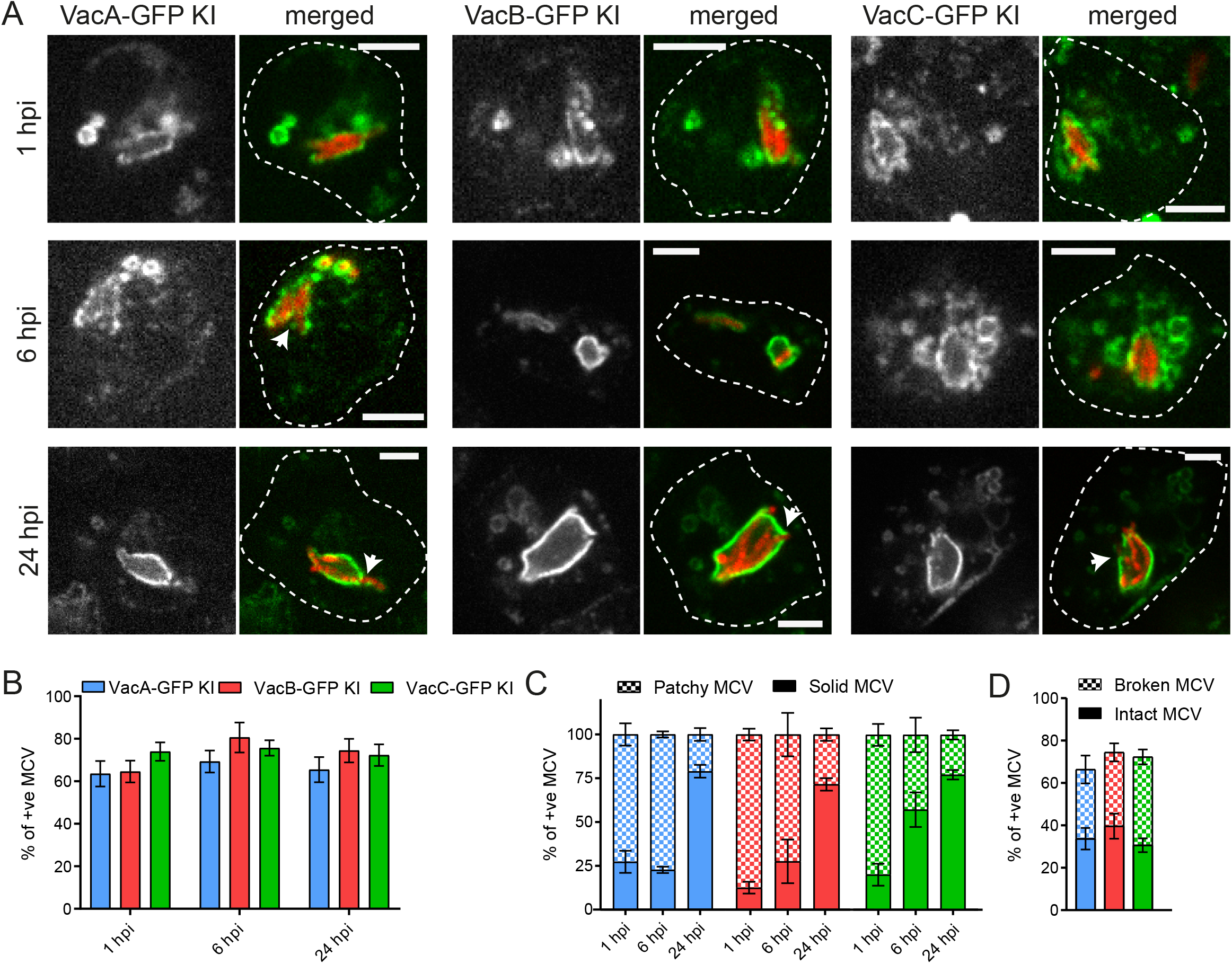
Vacuolins gradually accumulate on the MCV throughout the infection. **A.** Representative images of indicated Vac-GFP KI cell lines infected with wt *M*. *marinum* expressing mCherry. Arrows, broken MCVs; scale bar, 5 μm. **B.** Quantification of A. representing the proportion of MCVs positive for each vacuolin (mean ± s.e.m., N=4, n≥200 MCVs). **C.** Quantification of A. Of all vac-positive MCVs, the percentage of MCVs showing a patchy or solid vacuolin-coat were counted (mean ± s.e.m., N=4, n≥200 MCVs). **D.** Same data as in B. 24 hpi time point, MCVs were separated into visibly broken or intact MCVs.

### Absence of vacuolins confers resistance to *M*. *marinum* infection

To characterize the role of vacuolins in the biogenesis of the MCV, we analyzed the impact of vacuolin gene deletions on the infection course with the newly described vacuolin KO mutants (Bosmani et al., 2020). We have previously shown that, despite a faster postlysosomal maturation, vacuolin KO mutants are able to grow on, kill and digest different Gram-negative and -positive bacteria, as efficiently as wt cells (Bosmani et al., 2020). To test whether growth on and killing of mycobacteria were affected upon vacuolin KO, different dilutions of vacuolin KO mutants and wt cells were plated on mycobacterial lawns and their capacity to form phagocytic plaques, i.e. to grow and digest the bacteria, was assessed (**Fig. 3A and S2**). We observed in particular, that absence of one or more vacuolins facilitated growth on *M*. *marinum* wt, with KO cells growing 10-fold better than wt cells (**Fig. 3A**). On the other hand, vacuolin KO conferred only a slight advantage for growth on lawns of less pathogenic mycobacteria but with no correlation with the pathogenicity, as for instance KO cells grew better on *M*. *smegmatis* lawns rather than on *M*. *marinum* ΔRD1. To test whether absence of vacuolins impairs intracellular mycobacterial growth, we performed two different infection assays (**Fig. 3B and C**). Intracellular bacterial growth was measured either by luminescence using *lux*-expressing *M*. *marinum*, as previously described (Arafah et al., 2013), or by fluorescence using GFP-expressing *M*. *marinum*. Wt and vacuolin double or triple KO mutants (ΔBC, ΔABC) were infected and, after removing extracellular bacteria, luminescence was measured in a plate reader (**Fig. 3B**) or fluorescence by flow cytometry (**Fig. 3C**). Initial levels of infection were measured in each experiment to ensure a comparable starting point of infection for each cell line, despite the phagocytic defect observed in vacuolin KO cells (Bosmani et al., 2020). We observed that, in absence of two or all three vacuolins, wt *M*. *marinum* was not able to grow as efficiently as in wt cells, in particular after 24 hpi (**Fig. 3B-C**). After 72 hours of measurement, intracellular growth of *M*. *marinum* was reduced by at least 50% in vacuolin KO cells (**Fig. 3B-C**). In addition, infection experiments were performed with luminescent *M*. *marinum* ΔRD1 mutant, and, as mentioned earlier (**Fig. 3A**), absence of vacuolins had no impact on the growth of this mutant bacterium (**Fig. 3B**). Moreover, with time, the percentage of infected vacuolin KO cells decreased faster than for wt cells (**Fig. 3D**). Normally, the percentage of infected cells decreases as cells grow and multiply, thus diluting the number of infected cells. But, after 24 hpi, infection level of ΔBC cells was 75% lower than wt, despite similar cell growth (**Fig. 3E**), perhaps indicating an active “curing” of *M*. *marinum* infection. In conclusion, vacuolins are important susceptibility factors, as their depletion confers resistance to infection with *M*. *marinum*.

**Figure 3:**
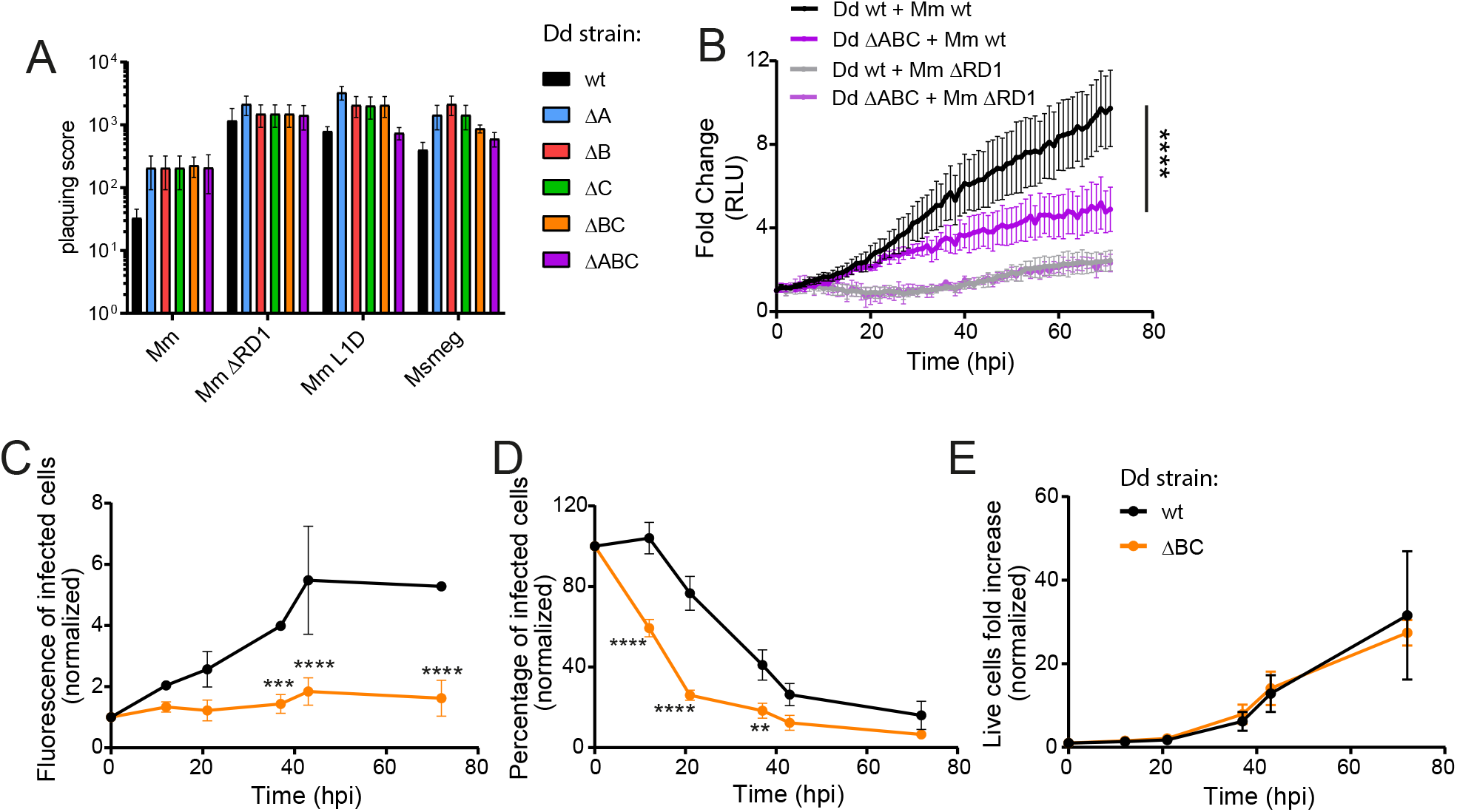
Absence of vacuolins confers resistance to infection. **A.** Different dilutions of wt or vacuolin KO cells were deposited on the indicated mycobacterial lawns mixed with *K. pneumoniae*. The plaquing score was determined using a logarithmic scale. Examples of plaques are shown in Fig. S2 (mean ± s.e.m., N=4). **B.** Wt and ΔABC cells were infected with luminescent *M*. *marinum* wt or ΔRD1 and luminescence measured every hour for 72 hours in a plate reader (mean fold change ± s.e.m. N=3, two-way ANOVA, ****p≤0.0001). **C-E.** Wt and ΔBC cells were infected with GFP-expressing *M*. *marinum*, cultured in shaking for 72 hours. At the indicated time points, fluorescence of infected cells (**C.**), percentage of infected cells (**D.**) and total number of cells (**E.**) was measured by flow cytometry (mean ± s.e.m., N=3, two-way ANOVA, *p≤0.05, **p≤0.01, ***p≤0.005, ****p≤0.0001).

### Vacuolins are not involved in manipulating acidification or proteolysis of the MCV

Resistance to infection can be attributed to different causes (**Fig. S3A-B**): (i) a more bactericidal compartment that may kill more efficiently the pathogen, (ii) premature escape from the MCV and subsequent capture by xenophagy, (iii) early release of the pathogen from the host cell, (iv) compromised access to nutrients and thus impaired growth, or (iv) incapacity to damage the MCV, and hence reduced escape to the nutrient-rich cytosol. To investigate at which step vacuolins are involved during infection, and how their absence may confer resistance, we infected wt and vacuolin KO cells with fluorescent *M*. *marinum* and characterized the MCV by immunofluorescence and time-lapse microscopy (**Fig. 4 and 5**).

**Figure 4.**
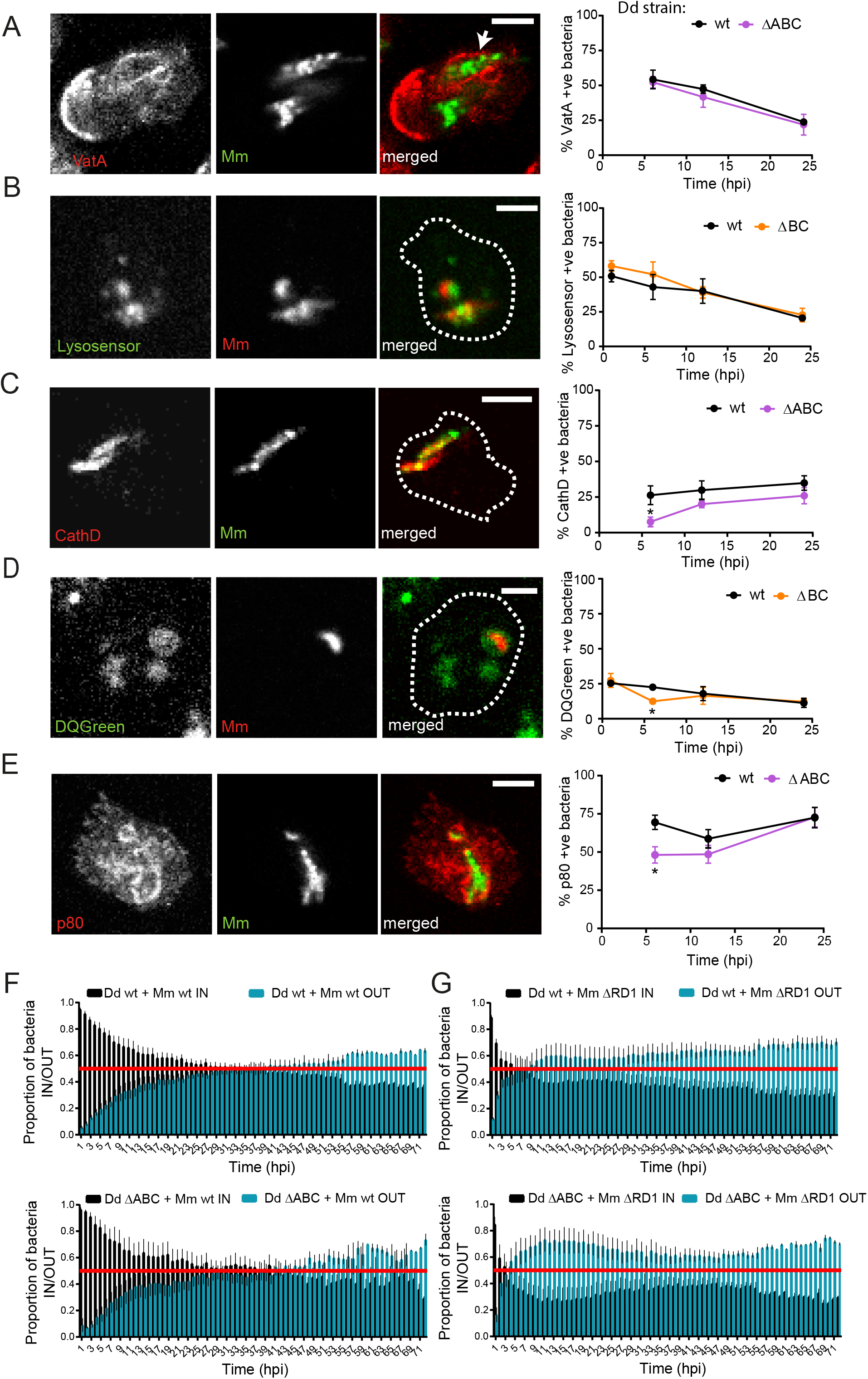
*M*. *marinum* does not reside in a more bactericidal compartment and is not released faster from vacuolin KO cells. **A, C and E.** Wt and ΔABC cells infected with GFP-expressing *M*. *marinum*, fixed at the indicated time points and immunostained with antibodies against VatA (**A.**), CathD (**C.**) or p80 (**E.**). Representative images of wt cells at 12 hpi are shown on the left. Scale bar, 5 μm. The proportion of marker-positive bacteria for each strain and time point is shown on the right. ((mean ± s.e.m., N=3, n≥150 MCVs). **B** and **D.** Wt and ΔBC cells infected with mCherry-expressing *M*. *marinum* imaged at the indicated time points after incubation with Lysosensor Green (**B.**) for 10 minutes or DQgreen (**D.**) for 1 hour before imaging. Representative images at 12 hpi are shown on the left. Scale bar, 5 μm. The proportion of marker-positive bacteria for each strain and time point is shown on the right ((mean ± s.e.m., N=3, n≥150 MCVs, *p≤0.05, two-way ANOVA). **F-G.** Wt and ΔABC cells infected with GFP-expressing *M*. *marinum* wt (**F.**) or ΔRD1 (**G.**) and imaged every hour by high content microscopy in the presence of antibiotics to inhibit extracellular growth. The proportion of intracellular and extracellular bacteria was analyzed by MetaXpress and plotted for each time point (mean ± s.e.m., N=3).

**Figure 5.**
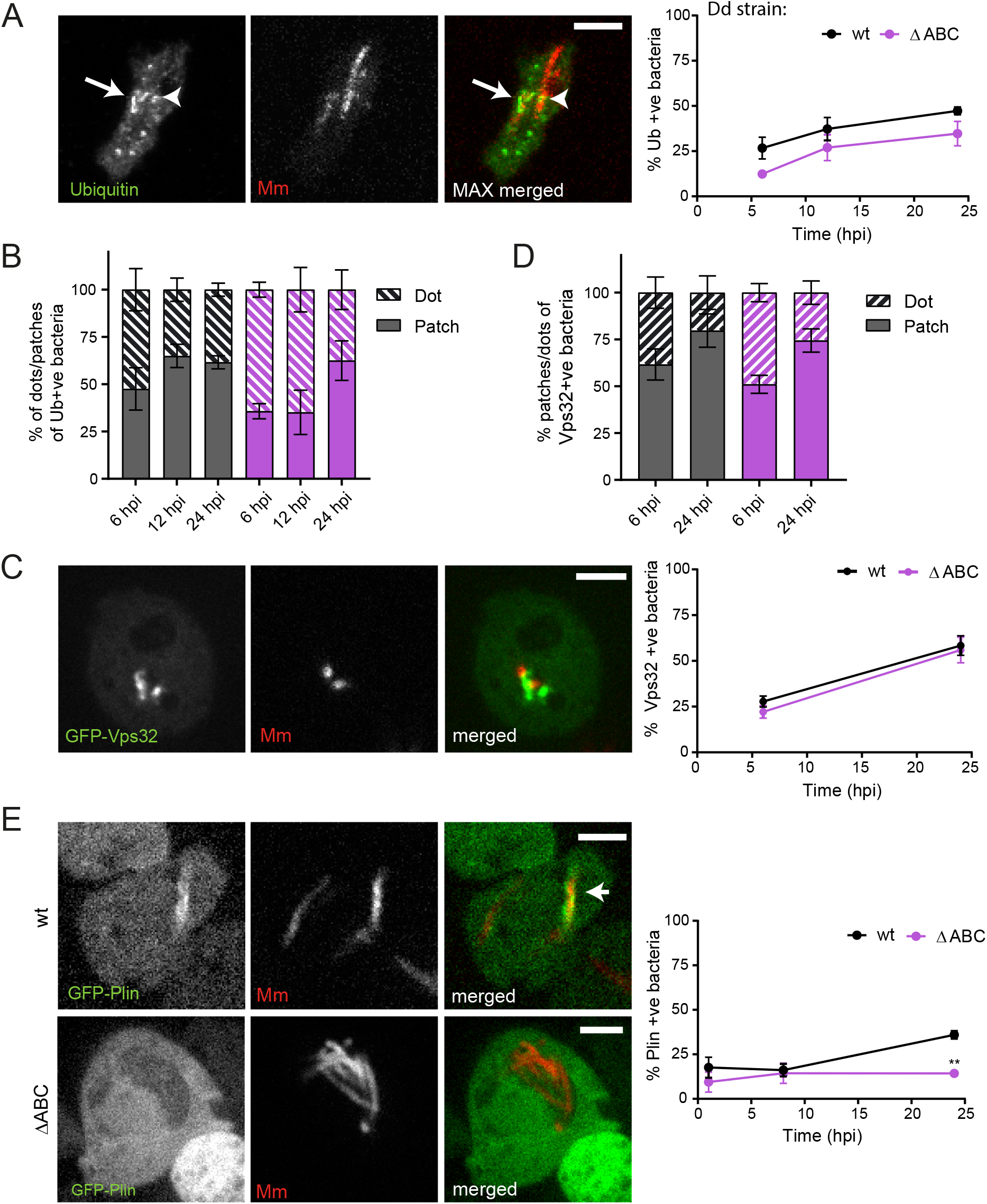
*M*. *marinum* cannot access the cytosol in vacuolin KO cells. **A.** Wt and ΔABC cells infected with mCherry-expressing *M*. *marinum* fixed at the indicated time points and immunostained with antibodies against Ubiquitin (FK2, mean ± s.e.m., N=3, n≥150 MCVs,). Representative images at 24 hpi are shown on the left. Scale bar, 5 μm. The proportion of marker-positive bacteria for each strain and time point is shown on the right. **B.** Same data as in A., of all Ub-positive MCVs, the percentage of MCVs showing a patchy or dotty Ubiquitin staining were counted (mean ± s.e.m.). **C-D.** Similar quantifications as in A-B., but of wt or ΔABC cells expressing GFP-Vps32 infected with mCherry-expressing *M*. *marinum* wt and imaged by time-lapse microscopy (N=2). **E.** wt or ΔABC cells expressing GFP-Perilipin (Plin) infected with mCherry-expressing *M*. *marinum* wt and imaged at the indicated time points by time-lapse microscopy (mean ± s.e.m., N=3, n≥150 MCVs, two-way ANOVA, **p≤0.01).

The v-ATPase is actively depleted from *M*. *tuberculosis* and *M*. *marinum*-containing compartments (Hagedorn and Soldati, 2007; Wong et al., 2011). In addition, proteolytic enzymes have been involved in resistance against intracellular pathogens, namely, *M*. *tuberculosis* regulates the expression of several cathepsins which otherwise would hinder its growth in macrophages (Pires et al., 2016). We first assayed whether absence of vacuolins might render the MCV more acidic (**Fig. 4-B**). The v-ATPase subunit VatA was present at about 50% of MCVs at 6 hpi, but was almost completely absent from MCVs after 24 hpi. No significant difference in v-ATPase content or depletion from the MCV was observed in wt compared with ΔABC cells (**Fig. 4A**). To measure acidity in the MCV itself, we incubated cells with the pH-dependent LysoSensor Green probe, which fluoresces in acidic compartments (**Fig. 4B**). In accordance with the previous result, the percentage of acidic compartments throughout the infection reflected well the number of VatA-positive MCVs and no difference was observed between wt and ΔABC cells. This indicates that, in absence of vacuolins, MCVs are not more acidic. To assay whether the MCV is more proteolytic, we monitored delivery of cathepsin D in the MCV (**Fig. 4C**) and measured proteolysis in the MCV using DQGreen-BSA, which is self-quenched but fluoresces upon proteolytic cleavage of BSA and release of DQGreen in the lumen (**Fig. 4D**). Throughout the infection, about 25 to 35% of *M*. *marinum* colocalized with CathD in wt cell. Interestingly, only 7% of the MCVs colocalized with CathD at 6 hpi in ΔABC cells compared to over 25% in wt cells, although at later time points CathD association in vacuolin KO mutants was similar to wt (**Fig. 4C**). Furthermore, as was previously shown (Cardenal-Muñoz et al., 2017), the *M*. *marinum*-containing compartment was poorly proteolytic, with no more than 25% DQGreen-positive compartments throughout the infection course (**Fig. 4D**). No main difference between wt and vacuolin KO cells was observed, except at 6 hpi, as for CathD. In conclusion, these results show that *M*. *marinum* is not exposed to a more acidic or proteolytic environment in absence of vacuolins. We have previously shown that vacuolin KOs have a slightly faster lysosome biogenesis, with CathD being delivered and/or retrieved faster from the phagosome during maturation (Bosmani et al., 2020). Given the differences in proteolysis and CathD association at 6 hpi, we tested whether another lysosomal-postlysosomal marker, p80, was differentially associated with the MCV in wt compared to vacuolin KO cells (**Fig. 4E**). Interestingly, at 6 hpi, p80 was less associated with the MCV in ΔABC cells (50% vs 70% in wt cells), although this was not the case at later time points. Overall, a notable difference in the association of lysosomal and postlysosomal markers was only observed at early time points, probably reflecting vacuolins’ role in lysosomal biogenesis. However, this phenotype is probably not sufficient to explain the resistance caused by absence of vacuolins, and thus we sought to further dissect this mechanism.

### Vacuolins are not involved in release of *M*. *marinum*

*M*. *marinum* can be released by the host cell by three mechanisms: cell death, exocytosis and a non-lytic actin-dependent egress termed ejection (Hagedorn et al., 2009; Cardenal-Muñoz et al., 2018). We reasoned that early release by any of these mechanisms may explain the lower percentage of infected cells, and thus the lower bacterial load, of vacuolin KO mutants. To test this, wt and ΔABC cells infected with GFP-expressing wt *M*. *marinum* or the ΔRD1 mutant, which is released by exocytosis more efficiently and earlier than wt bacteria (López-Jiménez et al., 2019), were imaged by high-content microscopy every hour for 72 hours (**Fig. 4F-G**). Cells were incubated during the whole experiment in medium containing antibiotics to prevent extracellular bacterial growth, and the proportion of extracellular versus intracellular bacteria was measured. Although we observed release from the host cell as early as 3 hpi on average in wt cells, 50% of all imaged bacteria were found in the extracellular space at 27 hpi (**Fig. 4F**). In ΔABC cells, bacteria were released in a similar dynamic and extent, suggesting that *M*. *marinum* is not released faster from vacuolin KO mutants. The ΔRD1 *M*. *marinum* mutant, as expected, was released much faster from wt cells than wt bacteria (**Fig. 4G**). In this case, 50% of all imaged bacteria were found in the extracellular space as early as 5 hpi. In ΔABC cells, the plateau was reached slightly faster, at 3 hpi. Nevertheless, these data suggest that early release is not a mechanism that fully explains the resistance to *M*. *marinum* wt infection in vacuolin KO mutants.

### Absence of vacuolins impairs *M*. *marinum* escape from the MCV

*M*. *marinum* resides inside the MCV and damages its membrane in an ESX-1-dependent manner, eventually resulting in escape to the cytosol (Cardenal-Muñoz et al., 2018; López-Jiménez et al., 2018). The ΔRD1 *M*. *marinum* mutant, which is strongly weakened in inducing damage and escaping from its compartment, is also strongly attenuated in *D*. *discoideum* cells (Cardenal-Muñoz et al., 2017). Similarly, in *D*. *discoideum* cells impaired in repairing the ESX-1-dependent damage, *M*. *marinum* escapes faster to the cytosol but is rapidly recaptured by autophagy, thus inhibiting its growth (López-Jiménez et al., 2018). Therefore, both earlier escape to the cytosol or impaired escape can result in reduced bacterial growth and we hypothesized that vacuolins modulate *M*. *marinum* escape from or //*retention inside the MCV. Ubiquitination of bacteria is one of the first signals of bacteria escape from the MCV and it triggers the recruitment of the autophagy machinery. In parallel, the ESCRT-III subunit Vps32 is found very early on damaged MCV membranes and mediates repair of membranes injuries inflicted by the ESX-1 secretion system (López-Jiménez et al., 2018). To dissect the mechanism by which vacuolins impair bacteria escape, we monitored ubiquitin and Vps32 recruitment at MCVs in wt and ΔABC cells (**Fig. 5A-D**). Ubiquitin was found to accumulate similarly in both cell lines, but to a lower extent (although not-significantly) in vacuolin KO mutants (**Fig. 5B**). Interestingly, the appearance of ubiquitin was quite different, especially in the first 12 hours of infection, with 65% of bacteria covered in ubiquitin patches in wt cells, and only 35% in ΔABC cells (**Fig. 5B**). On the other hand, GFP-Vps32 accumulated equally well in the vicinity of bacteria in wt compared to ΔABC cells (**Fig. 5C**), with only a small difference in the appearance of its recruitment (**Fig. 5D**). It was previously shown that the lipid droplet protein, perilipin (Plin), decorates bacteria once they gain access to the cytosol (Barisch et al., 2015), we thus monitored association of GFP-Plin with the bacteria in wt and ΔABC cells. (**Fig. 5E**). *M*. *marinum* was only poorly associated with Plin (15%) at 8 hpi in both wt cells and ΔABC, indicating that only a minor fraction of the bacteria is fully exposed to the cytosol at that stage, irrespective of the presence or absence of vacuolins. However, around 24 hpi, *M*. *marinum* gained access to the cytosol of wt cells and consequently was increasingly associated with GFP-Plin (45%). On the other hand, bacteria in ΔABC cells remained significantly less associated with GFP-Plin (15%). Together, these results suggest that in absence of vacuolins, *M*. *marinum* escapes less efficiently from the MCV, presumably by inflicting less membrane damage.

### ESAT-6-induced membrane damage requires vacuolins

*M*. *marinum* induces membrane damage via the ESX-1-mediated secretion of the small peptide ESAT-6, which inserts into host membranes (Gao et al., 2004; de Jonge et al., 2007). We hypothesized that ESAT-6 inserts in specific membrane domains containing vacuolins. To test this, we assayed *in vitro* whether recombinant Mtb ESAT-6 is able to bind host membranes in presence or absence of vacuolins (**Figs. 6A and S4A-B**). Host membranes of wt cells were purified and incubated with rESAT-6 at different pH and with different membrane-to-rESAT-6 ratios (**Fig. S4A-B**). In our conditions, the best binding to wt membranes was observed at pH 6. This result is in accordance with the fact that the pH measured directly inside the MCV in *D*. *discoideum*, using *M*. *marinum* coated with FITC and TRITC, is around 6 during most of the infection course, contrary to the marked acidification monitored around an avirulent *M*. *marinum* mutant (**Fig. S4C**). We then incubated rESAT-6 with membranes from wt and ΔABC cells in these specific conditions, and observed a reduction of about 50% of rESAT-6 association with membranes purified from vacuolin KO mutants (**Fig. 6A**). ESAT-6 is thought to prefer binding to ordered, cholesterol-rich membranes (de Jonge et al., 2007; Augenstreich et al., 2017). We wondered whether vacuolins are also part of these microdomains, in analogy to flotillins that are known lipid raft markers. To test this, Vac-GFP KI or overexpressing (OE) cells were lysed in cold Triton X-100 and Triton-insoluble fractions (TIF) were further floated by centrifugation in a sucrose gradient (**Fig. S5**). Surprisingly, and contrary to flotillins, only a small fraction of each vacuolin isoform was visible in the TIFs. Nevertheless, we hypothesized that vacuolins might associate more with ordered domains during infection (**Fig. 6B**). VacC-GFP KI cells were mock infected, or infected with wt *M*. *marinum*, and TSF and TIF/TIFF fractions were isolated. On average, VacC-GFP was found two-fold more enriched in TIF at 18 hpi than LmpB, a known marker of ordered domains in *D*. *discoideum* (Janssen et al., 2001). In conclusion, our results suggest that vacuolins, which are enriched in ordered domains during infection, may facilitate ESAT-6 insertion into host membranes, thus potentiating its membrane-damaging action.

**Figure 6.**
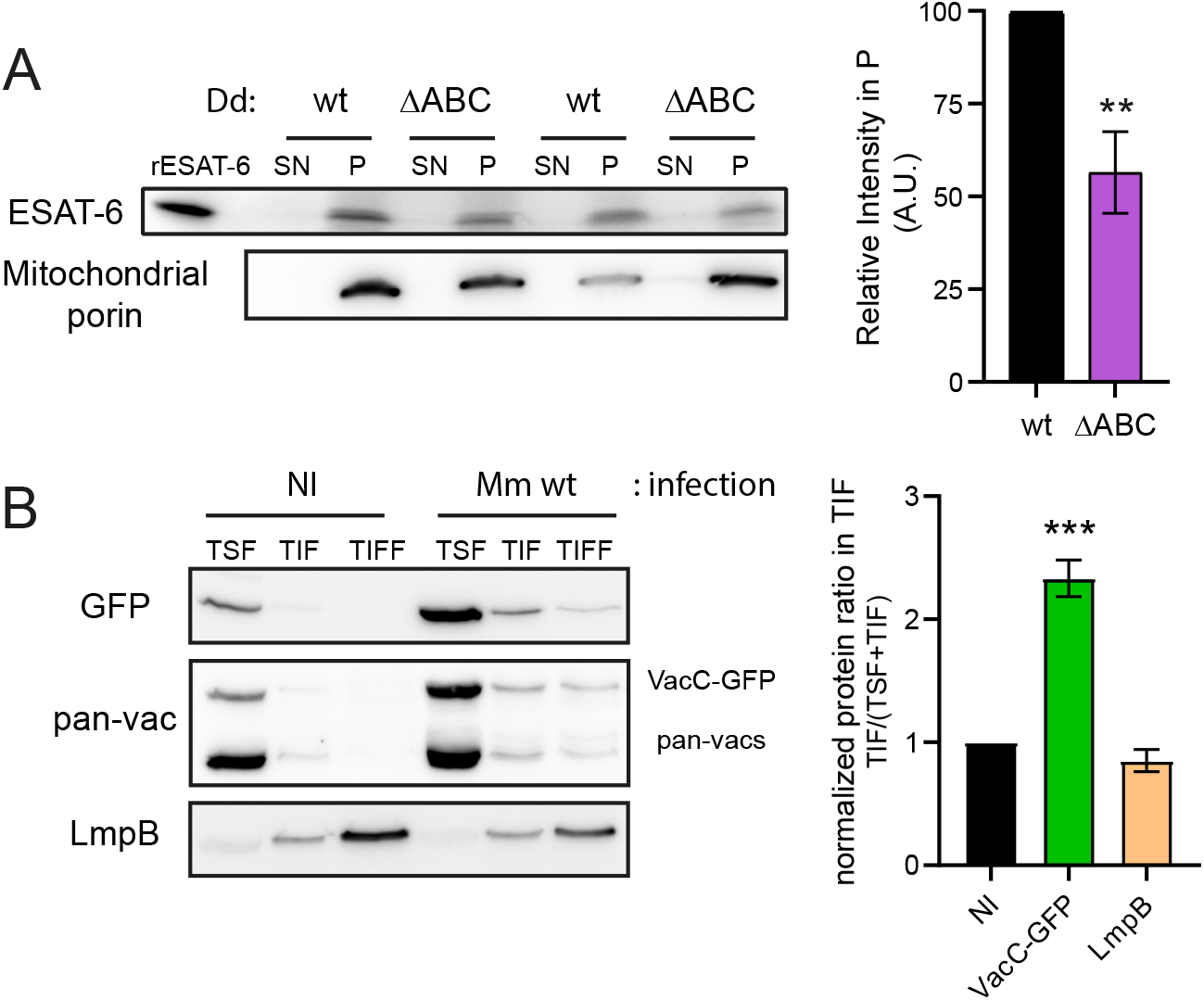
Microdomains enriched in vacuolins and cholesterol facilitate ESAT-6 membrane insertion. **A.** Recombinant ESAT6 (rESAT-6) was incubated with purified membranes of wt or ΔABC cells, then membranes were ultracentrifuged and separated into supernatant (SN) and pellet (P) fractions. Identical amounts were loaded for each fraction and immunoblotted with the indicated antibodies. Left: 2 independent replicates are shown. Right: quantification of ESAT-6 bands normalized to lane background and wt (mean ± s.e.m., N≥3, unpaired *t*-test, **p≤0.01). **B.** VacC-GFP KI cells were mock infected or infected with wt *M*. *marinum* and, after 16 hpi, lysed in cold Triton X-100. The Triton soluble (TSF) and insoluble (TIF) fractions were recovered, and the floating fraction (TIFF) after floatation on a sucrose gradient. Equal amounts of each fraction were loaded and immunoblotted with the indicated antibodies. Left: Representative image of 3 independent experiments. Right: quantification of the fraction of VacC found in the TIF compared to the total (TSF+TIF) (mean ± s.e.m., N=3, unpaired *t*-test, **p≤0.01.

## DISCUSSION

In *D*. *discoideum*, vacuolins are integral membrane proteins that oligomerize and thus define specific microdomains of the phagosomal membrane (Bosmani et al., 2020). Here, we propose that vacuolins are host factors that are highjacked by the pathogenic *M*. *marinum* and required for the efficient membranolytic activity of its secreted peptide ESAT-6 involved in membrane damage and escape from the MCV.

Our data show that vacuolins, in particular VacC, are highly induced, both at the mRNA and protein levels throughout the infection with wt *M*. *marinum*, but not with ΔRD1 or *M*. *smegmatis* (Fig. 1). We propose that VacC is mainly induced early on by mycobacteria species in general, perhaps resulting from recognition of specific mycobacterial pathogen associated molecular patterns (PAMPs). However, when the mycobacterium has a functional ESX-1 secretion system, induction of VacC is sustained, perhaps due to sensing of the damage caused by the secretion of membranolytic factors (Fig. 1C-F). Whether the induction of *vacC* is a host stress response or specifically and directly induced by pathogenic *M*. *marinum* remains to be determined. We also show that, while initially vacuolins are present at the MCV in patches, as the infection progresses, the bacterium becomes completely surrounded by a vacuolin-positive membrane (Fig. 2). Interestingly, MCVs exhibited a VacC-GFP positive coat earlier than with the other vacuolins. This could be a consequence of the higher expression of VacC during infection, or a higher tropism of this protein for the MCV.

We have previously described a role for VacB as a susceptibility factor involved in *M*. *marinum* infection (Hagedorn and Soldati, 2007). However, because the mutant used in that study was in fact a double vacuolin B and C KO (Bosmani et al., 2020), we present a more complete study and dissection of the role of all three vacuolins in the establishment of the *M*. *marinum* replicative niche. We confirm that absence of VacB, but also VacA or VacC, confers resistance to infection with *M*. *marinum* (Fig. 3). Interestingly, absence of vacuolin facilitates growth and plaque formation on a lawn of *M*. *marinum*, suggesting that vacuolin KO cells are more immune to the pathogenicity of *M*. *marinum* than wt cells (Fig. 3A). Importantly, we observe that growth on the ΔRD1 *M*. *marinum* mutant is as impaired in wt cells as in vacuolin KO mutants (Fig. 3A-B). In other words, absence of vacuolins seems to specifically impact growth of wt *M*. *marinum*, to a lesser extent that of *M*. *smegmatis*, but not of the ΔRD1 mutant. This corroborates the hypothesis that vacuolins are specifically hijacked by M. *marinum* and involved in establishment of the *M*. *marinum* niche in an ESX-1 and membrane damage-dependent manner.

We had previously shown that absence of VacB facilitates the accumulation of the v-ATPase at the MCV and hypothesized that VacB was involved in preventing association and/or assisting in recycling of the v-ATPase (Hagedorn and Soldati, 2007). In the present study, however, we find that the MCV of vacuolin KO mutants is not more proteolytic nor more acidic than in wt cells (Fig. 4 A-D). This indicates that vacuolins are not involved in preventing acidification of the MCV, nor in rendering the compartment less proteolytic. In addition, bacteria do not egress faster from vacuolin KO mutants (Fig. 4 E-F) but appear to reside longer in a more intact MCV (Fig. 5E). In fact, our data show that vacuolin KO cells are more resistant to infection because *M*. *marinum* is less efficient at escaping from its compartment. These results reinforce the idea that escape to the cytosol is required for optimal growth of *M*. *marinum* in its host and vacuolins play an important role in modulating mycobacteria escape.

How do vacuolins participate in *M*. *marinum* escape from the MCV? One hypothesis is that less damage would be produced at the MCV in absence of vacuolin microdomains. Two markers of damage and repair at the MCV are ubiquitin and the ESCRT-III subunit Vps32, respectively (López-Jiménez et al., 2018). We show that in absence of vacuolins, ubiquitination in the vicinity of the bacterium, which is one of the earliest signs of damage, is slightly lower, and less important in the first 12 hpi, suggesting that in absence of vacuolins damage is less extensive (Fig. 5A-B). However, surprisingly, this was not the case for Vps32 recruitment, as only a small non-significant change was observed between the patchiness and dotty morphology of the Vps32-positive areas (Fig. 5C-D). Therefore, our data might indicate that membrane repair is not affected by KO of vacuolins. On the other hand, because we measured the percentage of Vps32-positive bacteria, and since VacABC KO cells are by definition more resistant to infection, by 24 hpi, we are focusing on a subpopulation of cells that are still infected, while the majority of them are not anymore. However, taken together, our data about perilipin recruitment and ubiquitination lead us to conclude that in absence of vacuolins, bacteria escape less to the cytosol, probably as a consequence of inducing less membrane damage.

*M*. *marinum* and Mtb secrete ESAT-6, which has long been described as a pore-forming toxin (Gao et al., 2004; de Jonge et al., 2007; Leon et al., 2012; Augenstreich and Briken, 2020). ESAT-6 is secreted together with CFP-10, its putative chaperone, through the ESX-1 secretion system. In order to damage the membrane, ESAT-6 first needs to dissociate from CFP-10, which occurs at low pH, and was recently shown to require acetylations of ESAT-6 (de Jonge et al., 2007; Aguilera et al., 2020). Moreover, PDIMs (phthiocerol dimycocerosates), which are bacteria lipids, may activate ESAT-6 and participate in membrane damage (Augenstreich et al., 2017). During *M*. *marinum* infection of *D*. *discoideum* we did not monitor any significant drop in pH in the MCV for the average population of infected cells (Fig. S4C), probably due to early membrane damage caused by the bacterium leading to proton leakage. But our previously published results and those we present here, clearly document the fact that *M*. *marinum* experiences transient drops a of pH, resulting in a fraction of MCVs being LysoSensor-positive during the early stage of infection (Cardenal-Muñoz et al., 2017 and Fig. 4B). We assume that the drops in pH are transient, but are sufficient to trigger pH-dependent stimulation of genes such as Mag24-1 (Hagedorn and Soldati, 2007). As a consequence, the conditions that allow the dissociation of ESAT-6 from its chaperone are probably met. Alternatively, or in addition, we favour the idea that a host membrane component could facilitate the insertion of ESAT-6 in the MCV membrane. In that respect, it has been reported that ESAT-6 inserts into cholesterol-rich and ordered domains (de Jonge et al., 2007; Augenstreich et al., 2017), and we reason that vacuolins, which are functional homologues of flotillins, might be enriched in these domains. Indeed, and interestingly, while vacuolins were not enriched in Triton insoluble fractions at steady state in uninfected cells (Fig. S5), a significant increase in their association with these domains was observed after 16 hpi (Fig. 6B). This was reminiscent of our localization data (Fig. 2), which indicate that vacuolins are first present in small patches at the MCV but, as time progresses, they become highly enriched. This suggests that over time the lipid composition of the MCV undergoes significant changes, which may enable the bacterium to cause progressive and extensive damage that finally allows its escape. It is unclear, though, whether vacuolin enrichment is a cause or a consequence of this change.

Other pathogenic bacteria are known to require ordered domains in order to establish their infection. *Shigella flexneri* and *P. aeruginosa*, for example, co-opt the lipid rafts of their host membranes for their entry (Grassmé et al., 2003; Goot et al., 2004; Brandel et al., 2021). In addition, it was also proposed that these pathogens use lipid rafts to damage the compartment in which they reside. For instance, *S. flexneri* requires contact of its type three secretion system with the host lipid rafts to allow secretion of its virulence factors and thus induce membrane damage (Goot et al., 2004). Our data suggest that vacuolins, and possibly ordered domains, are necessary for the proper insertion of ESAT-6 into host membranes (Fig. 6A).

To conclude, here we propose that *M*. *marinum* controls the composition of the membrane of the MCV, and manipulates the host proteins of the vacuolin family, from their expression to their localization, to allow membrane damage, escape and survival inside the amoeba *D*. *discoideum*.

## MATERIALS AND METHODS

### *D*. *discoideum* strains, culture and plasmids

*D*. *discoideum* strains and plasmids are listed in Supplementary Table 1. Cells were axenically grown at 22°C in HL5c medium (Formedium) supplemented with 100 U/mL of penicillin and 100 μg/mL of streptomycin (Invitrogen). Plasmids were transfected into *D*. *discoideum* by electroporation and selected with the relevant antibiotic. Hygromycin was used at a concentration of 15 μg/mL, Blasticidin and G418 were used at a concentration of 5 μg/mL.

### Mycobacterial strains and culture

The mycobacterial strains used in this study are listed in Table S1. Mycobacteria were grown in Middlebrook 7H9 (Difco) supplemented with 10% OADC (Becton Dickinson), 0.2% glycerol and 0.05% Tween 80 (Sigma Aldrich) at 32°C in shaking culture at 150 r.p.m in the presence of 5 mm glass beads to prevent aggregation. Hygromycin was used at a concentration of 100 μg/mL (mCherry), kanamycin was used at a concentration of 50 μg/mL (GFP/DsRed expression) or 25 μg/mL (*lux* expression).

### Antibodies, reagents, western blotting and immunofluorescence

Recombinant nanobodies with the Fc portion of rabbit IgG, which recognize specifically VacA or VacB were previously characterized (Bosmani et al., 2020). The other following antibodies were used: pan-vacuolin (Dr. M. Maniak (Jenne et al., 1998)), VatA (Dr. M. Maniak (Jenne et al., 1998)), p80 (purchased from the Geneva Antibody Facility), cathepsin D (Dr. J. Garin (Journet et al., 1999)), ubiquitin FK2 (Enzo Life Sciences), and GFP (pAb from MBL Intl., mAb from Abmart). Goat anti-mouse or anti-rabbit IgG coupled to AlexaFluor 488, AlexaFluor 594, AlexaFluor 647 (Invitrogen) or to HRP (Brunschwig) were used as secondary antibodies.

After SDS-PAGE separation and transfer onto nitrocellulose membranes (Protran), immunodetection was performed as previously described (Schwarz et al., 2000) but with ECL Prime Blocking Reagent (Amersham Biosciences) instead of non-fat dry milk. Detection was performed with ECL Plus (Amersham Biosciences) using a Fusion Fx device (Vilber Lourmat). Quantification of band intensity was performed with ImageJ.

For immunofluorescence, infected *D*. *discoideum* cells were fixed with ultra-cold methanol (MeOH) at the indicated time points and immunostained as previously described (Hagedorn et al., 2006). Images were recorded with a Leica SP8 confocal microscope using a 63×1.4 NA oil immersion objectives.

### Time-lapse imaging

Infected cells were plated on a μ-dish (iBIDI) in filtered HL5c. After adherence, either 1 μm sections or time-lapse movies were taken with a spinning disc confocal system (Intelligent Imaging Innovations) mounted on an inverted microscope (Leica DMIRE2; Leica) using the 100×1.4 NA oil objective. Images were processed with ImageJ. Quantifications were performed manually. To stain acidic compartments, 1 μM LysoSensor Green DND-189 (ThermoFisher), a pH-dependent probe that becomes more fluorescent in acidic compartments, was added to the infected cells. After 10 min incubation, excess dye was washed off and cells were imaged for a maximum of 30 min. To stain compartments with proteolytic activity, 50 μg/mL DQ Green BSA (ThermoFisher), which releases fluorescent protein fragments upon proteolysis of the self-quenched BSA-associated Bodipy dye, was added to the infection sample one hour before imaging.

### Infection assays

Infections were performed as previously described (Hagedorn and Soldati, 2007; Arafah et al., 2013), with few modifications. After infection and phagocytosis, extracellular bacteria were washed off and attached infected cells were resuspended in filtered HL5c containing 5 μg/mL of streptomycin and 5 U/mL of penicillin to prevent growth of extracellular bacteria. Mock-infected cells were treated as above, but no bacteria were added.

To monitor both the host and pathogen during infection in a quantitative manner, cells were infected with GFP-expressing bacteria and analyzed by flow cytometry as previously described (Hagedorn and Soldati, 2007).

Growth of intracellular luminescent bacteria was measured as previously described (Arafah et al., 2013). Briefly, after infection with *luxABCDE*-expressing *M*. *marinum*, *D*. *discoideum* cells were counted and plated at different dilutions (from 1.3×10^5^ to 3×10^4^ cells/well) in a white F96 MicroWell plate (Nunc) covered with a gas permeable moisture barrier seal (Bioconcept). Luminescence was measured for 72 h with 1 h intervals with a Synergy Mx Monochromator-Based Multi-Mode Microplate Reader (Biotek) with constant 25°C.

To measure the proportion of intra and extracellular bacteria during the course of the infection, *D*. *discoideum* cells were infected, counted and plated in iBIDI 96-well μ-plates as described in Mottet et al., 2021.Infected cells were imaged for 72 h with 1 h intervals with a 40x objective with the ImageXpress Micro XL high-content microscope (Molecular Devices). Different parameters, including cell number, intracellular and extracellular bacterial number and fluorescence intensity, were extracted using the MetaXpress software (Molecular Devices, Mottet et al., 2021). The proportion of intracellular vs. extracellular bacteria was plotted by normalizing the fluorescence intensity of intra or extracellular bacteria to the total bacterial fluorescence at each time point.

To quantify the percentage of high versus low expression of Vac-GFP KI during infection, cells were plated at 24 hpi in iBIDI 96-well μ-plates and imaged with a 40x objective with the ImageXpress Micro XL high-content microscope. A cutoff of maximum intensity of 3000 was chosen to determine high and low expression based on the background intensity from non-infected cells. The proportion of high or low expression of each Vac-GFP was calculated for the population of non-infected cells (i.e., bystanders) and infected cells within the same well.

### Phagocytic plaque assay

Plaque formation of *D*. *discoideum* on a lawn of mycobacteria was monitored as previously described (Alibaud et al., 2011). 150-600 μl of a 5 × 10^8^ mycobacteria/mL culture were centrifuged and resuspended in 1.2 mL 7H9 containing a 1:10^5^ dilution of *K. pneumoniae* that had been grown overnight in LB. 50 μL of this suspension were deposited on wells from a 24-well plate containing 2 mL of 7H10-agar (without OADC). Serial dilutions of *D*. *discoideum* (10, 10^2^, 10^3^ or 10^4^ cells) were plated onto the bacterial lawn and plaque formation was monitored after 4-7 days at 25°C. To quantify cell growth on bacteria, a logarithmic growth score was assigned as follows: plaque formation up to a dilution of 10 cells received a score of 1000; when cells were not able to grow at lower dilutions, they obtained the corresponding lower scores of 100, 10 and 1.

### Quantitative real-time PCR (qPCR)

RNA from mock-infected cells or cells infected with *M*. *marinum* wt, ΔRD1 or *M*. *smegmatis* was extracted at the indicated time points using the Direct-zol RNA MiniPrep kit (Zymo Research) following manufacturer’s instructions. 1 μg of RNA was retro-transcribed using the iScript cDNA Synthesis Kit and polydT primers (Biorad). The cDNA was amplified using the primers listed in S2 Table and the SsoAdvanced universal SYBR Green supermix (Biorad). Amplimers for *vacA, vacB, vacC* and *gapdh* were detected on a CFX Connect Real-Time PCR Detection System (Biorad). The housekeeping gene *gapdh* was used for normalization. PCR amplification was followed by a DNA melting curve analysis to confirm the presence of a single amplicon. Relative mRNA levels (2^−ΔΔCt^) were determined by comparing first the PCR cycle thresholds (Ct) for the gene of interest and *gapdh* (ΔC), and second Ct values in infected cells vs mock-infected cells (ΔΔC).

### RNAseq

Following infection with GFP-expressing bacteria, infected and mock-infected cells were pelleted and resuspended in 500 μl of HL5c, passed through 30 μm filters and sorted by FACS (Beckman Coulter MoFlo Astrios). The gating was based on cell diameter (forward-scatter) and granularity (side-scatter). Of those, infected (GFP-positive) and non-infected (GFP-negative) sub-fractions were based on GFP intensity (FITC channel). Typically, ∼5 × 10^5^ cells of each fraction were collected for RNA isolation. RNA was isolated as above. Quality of RNA, libraries, sequencing and bioinformatic analysis were performed as previously described (Hanna et al., 2019).

### Cytosol-membrane separation and rESAT-6 incubation

10^9^ *D*. *discoideum* cells were washed in Sorensen-Sorbitol and resuspended in HESES buffer (HEPES 20 mM, 250 mM Sucrose, MgCl_2_ 5 mM, ATP 5 mM) supplemented with proteases inhibitors (cOmplete EDTA-free, Roche). Cells were homogenized in a ball homogenizer with 10 μm clearance. The post-nuclear supernatant was diluted in HESES buffer and centrifuged at 35’000 rpm in a Sw60 Ti rotor (Beckmann) for 1 hour at 4°C. The cytosol (supernatant) and membrane (MB, pellet) fractions were recovered. The protein concentration of the cytosol fraction was quantified by Bradford. Different quantities of membranes were tested (Fig S4), finally 400 μg of membranes were incubated with 12 μg of recombinant ESAT-6 (rESAT-6, BEI Resources, NR-49424) in HESES Buffer at pH 6, for 20 min at RT on a wheel. After ultracentrifugation at 45’000 rpm for 1 hour, 4°C, in a TLS-55 rotor (Beckmann), membranes were separated into supernatant (SN) and pellet (P) fractions. Equal amounts of SN and P were loaded for western blotting.

### Detergent resistant membrane isolation

10^8^ *D*. *discoideum* cells were washed in Sorensen-Sorbitol and resuspended in 1 ml of cold Lysis Buffer (Tris-HCl pH 7.5 50 mM, NaCl 150 mM, Sucrose 50 mM, EDTA 5 mM, ATP 5 mM, DTT 1 mM) with 1% Triton X-100 supplemented with proteases inhibitors. The lysate was then incubated at 4°C on a rotating wheel for 30 min. After centrifugation (5 min, 13’000 rpm, table-top centrifuge, 4°C), the supernatant (Triton Soluble Fraction, TSF) was collected and the pellet (Triton Insoluble Fraction, TIF) was resuspended in 200 μl of cold Lysis buffer without Triton X-100. The TIF was mixed with 800 μl of 80% sucrose (final concentration 65% sucrose), deposited at the bottom of an ultracentrifuge tube and overlayed with 2 ml of 50% sucrose and 1 ml of 10% sucrose. The TIF was then centrifuged at 55’000 rpm in a Sw60 Ti rotor (Beckmann) for 2 h at 4°C. The Triton Insoluble Floating Fraction (TIFF) was collected, acetone precipitated and resuspended in Laemmli Buffer. Equivalent amounts of each fraction (TSF, TIF and TIFF) were loaded for western blotting.

## ACKNOWLEDGEMENTS

We gratefully acknowledge Dr. P. Cosson (University of Geneva) and Dr. J. King (University of Sheffield) for discussions and suggestions and the staff of the Bioimaging Center for Microscopy, the FACS core facility and the IGE3 Genomics Platform at the Faculty of Sciences and Faculty of Medicine of the University of Geneva for their precious help. We thank Dr. D. Moreau and the ACCESS Geneva Imaging Facility of the University of Geneva for help with the high content microscopy experiments. This work was supported by Swiss National Science Foundation (SNF) grants 310030_169386 and 310030_188813 TS is a member of iGE3 (http://www.ige3.unige.ch).

## FIGURE LEGENDS

**Figure S1. Endogenous VacA and VacB are present on the MCV.** Representative images of wt cells infected with wt GFP-expressing *M*. *marinum*, fixed at indicated time points and immunostained with antibodies against endogenous VacA or VacB. Scale bar, 5 μm.

**Figure S2. Plaque assays with vacuolin KO cells.** Representative images of plaque assays of Fig. 3A taken at day 5.

**Figure S3.** Model depicting different possible explanations for a resistance to infection. **A.** The normal infection course of *M*. *marinum* in wt cells. **B.** Five different mechanisms could explain the resistance of ΔABC cells (see main text).

**Figure S4. r-ESAT-6 binds membranes at pH 6, the pH measured in the MCV lumen. A-B.** Recombinant ESAT6 (rESAT-6) was incubated with purified membranes of wt cells at different pH (A.) or ratios (B.), then membranes were ultracentrifuged and separated into supernatant (SN) and pellet (P) fractions. Identical amounts were loaded for each fraction and immunoblotted with the indicated antibody. **C.** Wt cells were spinoculated with *M*. *marinum* or *M*. *marinum*-L1D labelled with the fluorophores FITC (pH sensitive) and TRITC (pH insensitive). The ratio of both fluorophores was measured by plate reader (mean ± s.e.m., N=2, n≥3).

**Figure S5. r-ESAT-6 binds membranes at pH 6, the pH measured in the MCV lumen. A-B.** Vac-GFP KI (A) or Vac-GFP overexpressing (OE, B) cells were lysed in cold Triton X-100. The Triton soluble (TSF) and insoluble (TIF) fractions were recovered, and the floating fraction (TIFF) after floatation on a sucrose gradient. Equal amounts of each fraction were loaded and immunoblotted with the indicated antibodies. Representative images of 2 (A) or 3 (B) independent experiments.

