## Supplemental data for "Disruption of vacuolin microdomains in the host *Dictyostelium discoideum* increases resistance to *Mycobacterium marinum*-induced membrane damage and infection"

**Sup. Table 1.** List of *D. discoideum* strains used in this study.

| Strain/Plasmid | Relevant characteristics | Source/Reference |
|---|---|---|
| *D. discoideum* | | |
| Ax2(Ka) | Wild-type | |
| Ax2(Ka) ΔVacA knock-out strain | | (Bosmani et al., 2020) |
| Ax2(Ka) ΔVacB knock-out strain | | (Bosmani et al., 2020) |
| Ax2(Ka) ΔVacC knock-out strain | | (Bosmani et al., 2020) |
| Ax2(Ka) ΔVacBΔVacC knock-out strain | | (Bosmani et al., 2020) |
| Ax2(Ka) ΔVacAΔVacBΔVacC knock-out strain | | (Bosmani et al., 2020) |
| Ax2(Ka) VacA-GFP knock-in strain | | (Bosmani et al., 2020) |
| Ax2(Ka) VacB-GFP knock-in strain | | (Bosmani et al., 2020) |
| Ax2(Ka) VacC-GFP knock-in strain | | (Bosmani et al., 2020) |
| Mycobacteria strains | | |
| *M. marinum* M strain | Wild-type | L. Ramakrishnan (Washington University) |
| *M. marinum* ΔRD1 | | L. Ramakrishnan (Washington University) |
| *M. marinum* L1D | | L. Ramakrishnan (Washington University) |
| *M. smegmatis* | | G. Griffiths (EMBL, Heidelberg, Germany) |
| *D. discoideum* plasmids | | |
| pDNeoGFP-Plin | | (Du et al., 2013) |
| GFP-Vps32 | | (López-Jiménez et al., 2018) |
| Mycobacteria plasmids | | |
| pCherry10 | mCherry under control of the G13 promoter, $Hyg^r$ | (Carroll et al., 2010) |
| pMSP12::DsRed/GFP | DsRed/GFP under control of the MSP promoter, $Kan^r$ | (Cosma et al., 2004) |
| pMV306::*lux* | bacterial luciferase under control of the G13 promoter, $Kan^r$ | (Andreu et al., 2010) |

**Sup. Table 2.** List of oligos used in this study, F: forward, R: reverse

| **Oligonucleotide/Use** | **Sequence 5'-3'** |
|---|---|
| qPCR | |
| gapdhF | GGTTGTCCCAATTGGTATTAATGG |
| gapdhR | CCGTGGGTTGAATCATATTTGAAC |
| vacAF | CATTGGCAGATAACAAATCAGCATTGG |
| vacAR | ATCATTGGTGGCTTGACCTGGTTTAAT |
| vacBF | GAAATTAGATATCTTGGCTCAGTTGCAAA |
| vacBR | ATTACCACTTTCACTAGTATCTTCAACCAT |
| vacCF | TTAAAGTTGTCGAACAAATAGGTAGTC |
| vacCR | CCATCTTTTGGTGTTTGAATAATTTCAAG |

Bosmani et. al.,
Figure S1

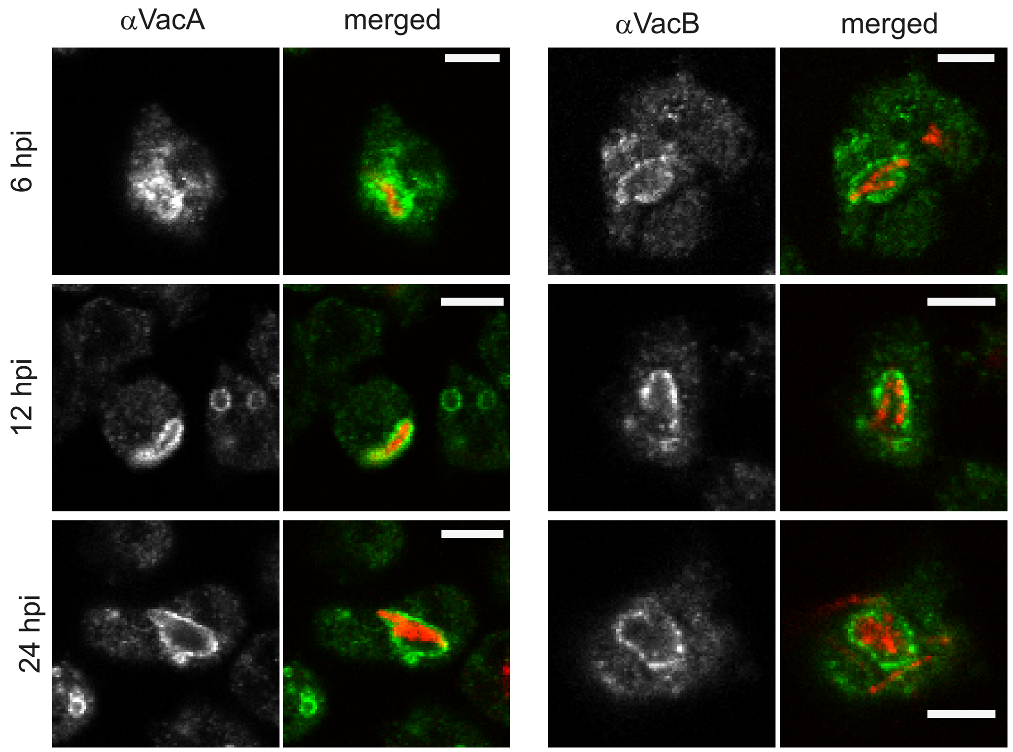

Bosmani et. al.,
Figure S2

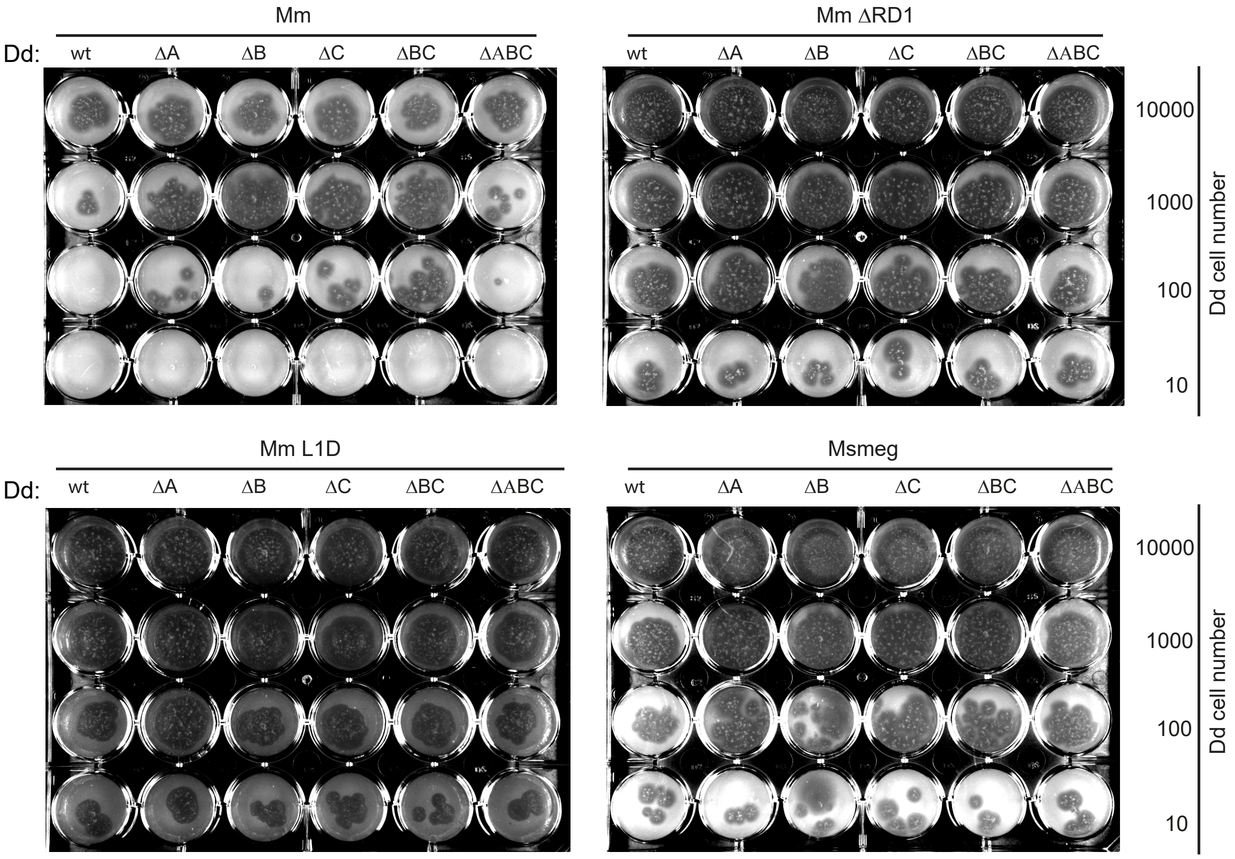

Bosmani et. al.,
Figure S3

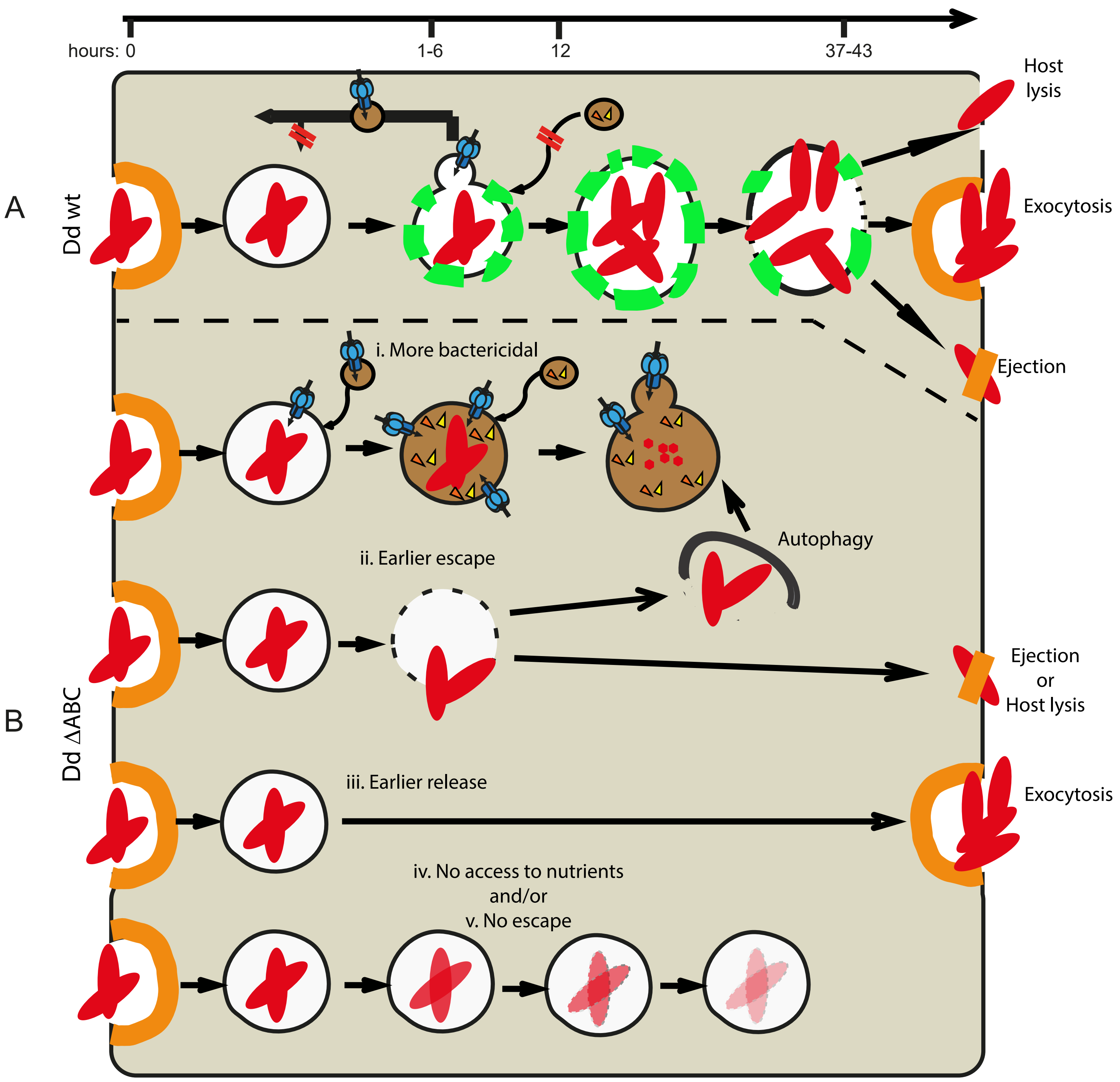

Bosmani et. al.,
Figure S4

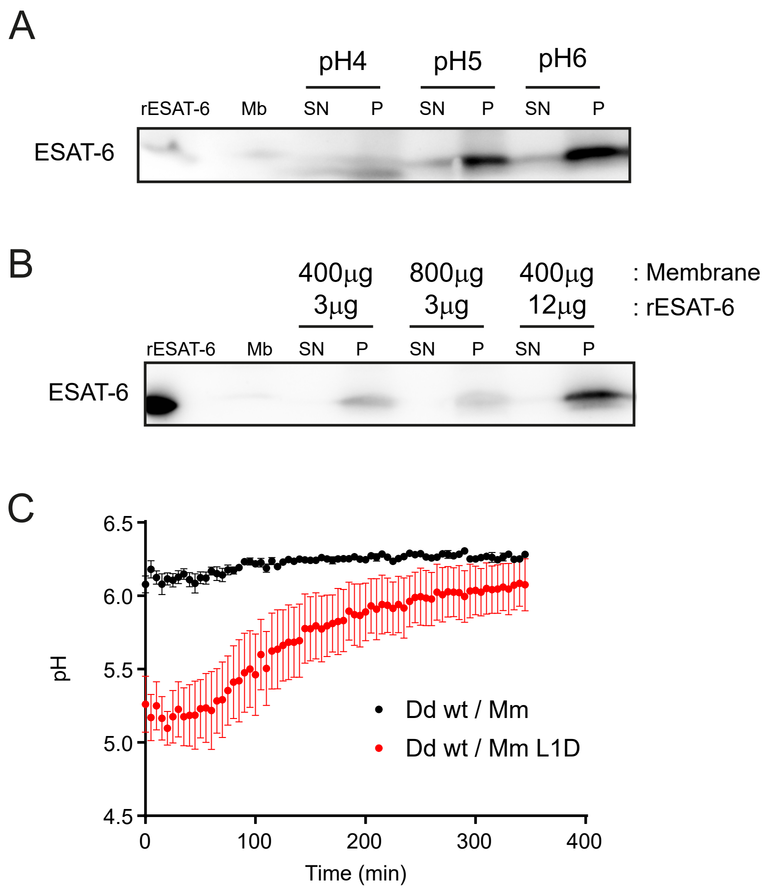

Bosmani et. al.,
Figure S5

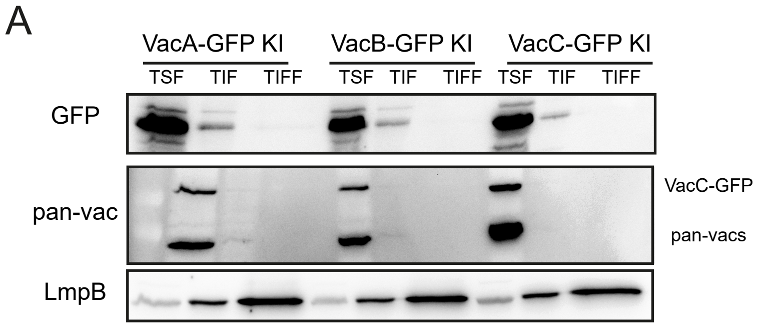


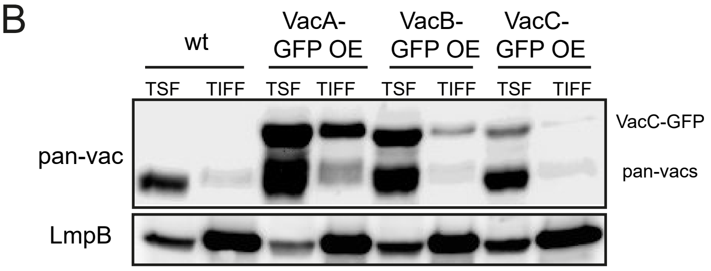
